# Hypothalamic Overexpression of Neurosecretory Protein GL Leads to Obesity in Mice

**DOI:** 10.1101/2021.03.01.433475

**Authors:** Yuki Narimatsu, Keisuke Fukumura, Kenshiro Shikano, Eiko Iwakoshi-Ukena, Megumi Furumitsu, George E. Bentley, Lance J. Kriegsfeld, Kazuyoshi Ukena

## Abstract

**Introduction:** The mechanisms underlying obesity are not fully understood, necessitating the creation of novel animal models for investigation of metabolic disorders from the cellular-molecular to behavioral levels of analysis. We have previously found that neurosecretory protein GL (NPGL), a newly-identified hypothalamic neuropeptide, is involved in feeding behavior and fat accumulation in rats. Given the broad availability of genetic tools in mice, the present investigation sought to establish a mouse model of NPGL-induced obesity.

**Methods:** We overexpressed the NPGL-precursor gene (*Npgl*) in the hypothalamus using adeno-associated virus in C57BL/6J mice fed normal chow (NC) or a high-calorie diet (HCD). After 9 weeks of *Npgl* overexpression, we measured adipose tissues, muscle, and several organ masses in addition to food intake and body mass. To assess the effects of *Npgl* overexpression on peripheral tissues, we analyzed mRNA expression of lipid metabolism-related genes by quantitative RT-PCR.

**Results:** *Npgl* overexpression increased food intake, body mass, adipose tissues and liver masses, food efficiency, and circulating insulin levels under both NC and HCD, resulting in obesity observable within 8 weeks. Furthermore, we observed fat accumulation in adipose tissues and liver. Additionally, mRNA expression of lipid metabolism-related factors was increased in white adipose tissue and the liver after *Npgl* overexpression.

**Conclusion:** Taken together, the present study suggests that NPGL is an endogenous obesogenic factor that acts within a short period of time in mice. As a result, this animal model can be widely applied to study the etiology of obesity from genes to behavior.

## Introduction

The number of obese patients worldwide has recently reached 700 million [1]. Obesity is associated with many serious diseases, such as type 2 diabetes mellitus, hypertension, and hyperlipidemia [2–5]. The imbalance between energy intake and energy expenditure causes fat accumulation and eventually obesity [6]. Therefore, further research on energy homeostasis is required to prevent and treat metabolic disease. Energy homeostasis is regulated by factors produced by the brain and peripheral tissues. The neuropeptides α-melanocyte-stimulating hormone (α-MSH)—derived from proopiomelanocortin (POMC)—and neuropeptide Y (NPY) are known to be anorexigenic and orexigenic respectively. Both are produced in the hypothalamic arcuate nucleus (Arc) [7]. Agouti-related peptide (AgRP) is also produced by NPY-expressing neurons and inhibits the effect of α-MSH by antagonistic binding to melanocortin receptor type 4 (MC4R) [8]. Ghrelin and leptin are other well-known energy-related metabolic factors secreted by peripheral tissues. Ghrelin is an orexigenic factor secreted by the stomach, and it promotes feeding behavior by activating NPY/AgRP neurons [9]. Leptin is an anorexigenic hormone released from white adipose tissue (WAT) [10, 11]. Moreover, insulin accelerates fat accumulation by inhibiting lipolysis [12]. It is clear that many factors are involved in the regulation of energy intake, but this complex system is still not fully understood. Thus, the identification and functional analysis of novel factors is required for understanding whole body energy homeostasis.

We recently found a novel gene related to energy homeostasis in chickens, rats, mice, and humans [13–15]. The small secretory protein produced by the novel gene was named neurosecretory protein GL (NPGL) because the C-terminus amino acid sequence is Gly-Leu-NH_2_ [13]. Analysis of the genome database revealed that NPGL is evolutionarily conserved throughout vertebrates [13]. To date, we have mainly investigated the functions and features of NPGL in rats and mice. Using morphological analysis, we found that NPGL is produced in the lateroposterior region of the Arc in mice, and NPGL-containing fibers project to several regions within the hypothalamus [15].

We recently showed that NPGL regulates feeding and fat accumulation in rodents. In rats, a 13-day chronic intracerebroventricular (i.c.v.) infusion of NPGL increased the WAT mass without changing body mass and modestly stimulated food intake under a high-calorie diet (HCD) [14]. Analysis of adeno-associated virus (AAV)-driven *Npgl* overexpression in rats for 6 weeks had more marked effects on feeding behavior and lipogenesis than the 13-day chronic infusion [14]. Similarly, 13-day chronic i.c.v. infusion of NPGL stimulated feeding behavior and mass gain due to fat accumulation in mice fed a HCD [16]. However, the induction of abnormally high body mass was not a characteristic result of NPGL treatment in these studies, leaving the possibility that NPGL induces obesity as an open question. In general, creation of diet-induced obesity (DIO) in rodents is costly and time-consuming, typically requiring 16–20 weeks [17–19]. Therefore, the discovery of a more efficient way to generate an animal model of obesity would be useful for studies aiming to elucidate the mechanism of energy metabolism regulation.

The aim of the present study was to determine the impact of NPGL on obesity in a widely-used animal model with broad foundational data and molecular-genetic tools available. We found that overexpression of the NPGL-precursor gene (*Npgl*) in the mouse hypothalamus elicited obesity within 8 weeks under a HCD. The present findings report the effects of *Npgl* overexpression on food intake, body mass, body composition, blood parameters, and the expression of lipid metabolism-related factors in mice fed normal chow (NC) or a HCD during the development of obesity.

## Materials and Methods

### Animals

Male C57BL/6J mice (7 weeks old) were purchased from Nihon SLC (Hamamatsu, Japan) and housed under standard conditions (25 ± 1°C under a 12-h light/12-h dark cycle) with *ad libitum* access to water and NC (CE-2; CLEA Japan, Tokyo, Japan) or a HCD (32% of calories from fat/20% of calories from sucrose, D14050401; Research Diets, New Brunswick, NJ). Animal surgery was performed under isoflurane anesthesia.

### Production of AAV-based Vectors

We followed a previously reported method to generate the overexpression AAV [14]. The full-length open reading frame of mouse *Npgl* was amplified from cDNA of the mediobasal hypothalamus and inserted into the pAAV-IRES-GFP expression vector (Cell Biolabs, San Diego, CA). The primers for mouse NPGL were 5’-CGATCGATACCATGGCTGATCCTGGGC-3’ for the sense primer and 5’-CGGAATTCTTATTTTCTCTTTACTTCCAGC-3’ for the antisense primer.

AAV-based vectors AAV-DJ/8-NPGL-IRES-GFP for NPGL (AAV-NPGL) and AAV-DJ/8-IRES-GFP for control (AAV-CTL) as shown in supplementary Fig. S1a were produced in 293AAV cells (Cat# AAV-100; Cell Biolabs) using the AAV-DJ/8 Helper Free Packaging System containing pAAV-DJ/8 and pHelper plasmids (Cell Biolabs). The triple plasmids (AAV-DJ/8-NPGL-IRES-GFP or AAV-DJ/8-IRES-GFP, pAAV-DJ/8, and pHelper) were mixed with the polyethylenimine MAX transfection reagent (PEI-MAX; Polysciences, Warrington, PA). The mixture was diluted with Opti-MEM I medium (Life Technologies, Carlsbad, CA) and added to 293AAV cells in 150-mm cell culture dishes. Transfected cells were cultured in DMEM containing 10% fetal bovine serum.

For the purification of AAV-based vectors, 3 days after transfection, the cells and supernatants were harvested and purified using chloroform and were condensed using Amicon Ultra-4 Centrifugal Filter Devices (100K MWCO; Merck Millipore, Billerica, MA).

For AAV titration, 1 μL of AAV solution was treated with RQ1 DNase (Promega, Madison, WI) according to the manufacturer’s instructions. Virus titers were determined by quantitative PCR with EGFP primer pairs. The primers for EGFP were 5’-ACCACTACCTGAGCACCCAGTC-3’ for the sense primer and 5’-GTCCATGCCGAGAGTGATCC-3’ for the antisense primer. After titration, the AAV-based vectors were prepared at a concentration of 1 × 10^9^ particles/µL and stored at −80°C until use.

### Npgl Overexpression

Mice were divided into two groups according to each diet (NC or HCD). For *Npgl* overexpression, mice were bilaterally injected with 0.5 µL/site (5.0 × 10^8^ particles/site) of AAV-based vectors (AAV-NPGL or AAV-CTL) using a Neuros Syringe (7001 KH; Hamilton, Reno, NV) into the mediobasal hypothalamic region with the coordinates 2.2 mm caudal to the bregma, 0.25 mm lateral to the midline, and 5.8 mm ventral to the skull surface.

*Npgl* overexpression was maintained for 9 weeks in mice fed NC or a HCD. *Npgl* overexpression was confirmed by quantitative RT-PCR at the endpoint. Food intake and body mass were measured weekly. Food efficiency (g/kcal) was calculated as body mass gain (g)/ cumulative food intake (kcal) [20]. Body composition and serum parameters were measured at the end of *Npgl* overexpression.

### Quantitative RT-PCR

The inguinal WAT (iWAT) and liver were dissected from mice and snap frozen in liquid nitrogen for RNA processing. Total RNA was extracted using QIAzol lysis reagent for the iWAT (QIAGEN, Venlo, Netherlands) or TRIzol reagent for the liver (Life Technologies) in accordance with the manufacturer’s instructions. First-strand cDNA was synthesized from total RNA using a ReverTra Ace kit (TOYOBO, Osaka, Japan).

The abbreviations for genes and sequences of primers used in this study are listed in Table 1 and Table 2, respectively. PCR amplifications were conducted with THUNDERBIRD SYBR qPCR Mix (TOYOBO) using the following conditions: 95°C for 20 s, followed by 40 cycles each consisting of 95°C for 3 s, and 60°C for 30 s. The PCR products in each cycle were monitored using a Bio-Rad CFX Connect (Bio-Rad Laboratories, Hercules, CA). Relative quantification of each gene was determined by the 2^−ΔΔCt^ method using ribosomal protein S18 (*Rps18*) for the iWAT, or β-actin (*Actb*) for the liver as an internal control [21].

**Table 1.** Abbreviation

| Gene | Name |
| --- | --- |
| *Acc* | Acetyl-CoA carboxylase |
| *Fas* | Fatty acid synthase |
| *Scd1* | Stearoyl-CoA desaturase 1 |
| *Gpat1* | Glycerol-3-phosphate acyltransferase 1 |
| *Chrebpα* | Carbohydrate-responsive element-binding protein $\alpha$ |
| *Cpt1a* | Carnitine palmitoyl transferase 1a |
| *Atgl* | Adipose triglyceride lipase |
| *Hsl* | Hormone-sensitive lipase |
| *Fgf21* | Fibroblast growth factor 21 |
| *Gapdh* | Glyceraldehyde 3-phosphate dehydrogenase |
| *Slc2a2* | Solute carrier family 2 member 2 |
| *Slc2a4* | Solute carrier family 2 member 4 |
| *Cd36* | Cluster of differentiation 36 |
| *Ppara* | Peroxisome proliferator-activated receptor $\alpha$ |
| *Pparγ* | Peroxisome proliferator-activated receptor $\gamma$ |
| *Pgc1α* | Peroxisome proliferator-activated receptor $\gamma$ coactivator 1α |
| *Tnfa* | Tumor necrosis factor $\alpha$ |
| *Adipoq* | Adiponectin |
| *Rps18* | Ribosomal protein S18 |
| *Actb* | β-Actin |

**Table 2.**
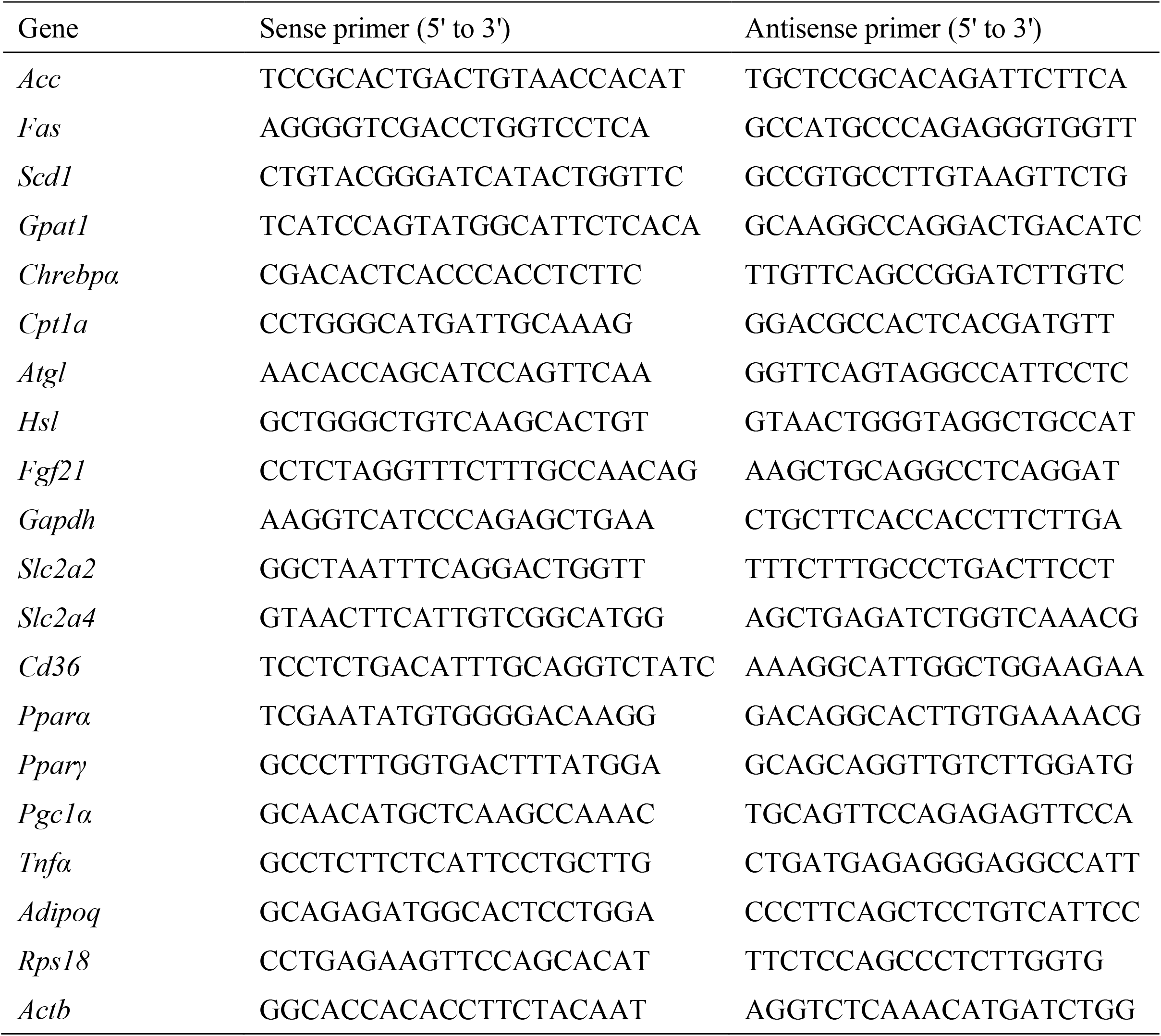
Sequences of oligonucleotide primers for quantitative RT-PCR

### Fatty Acid Analysis

For analysis of endogenous stearoyl-CoA desaturase 1 (SCD1) activity in the iWAT, lipids were extracted according to a previously described method [22]. The iWAT (50 mg) was extracted with 500 µL of chloroform:methanol (2:1) using a bead crusher (μT-12; TAITEC, Saitama, Japan), and 125 µL of distilled water was added and mixed by inversion. After incubation for 30 min, the sample was centrifuged at 3000 × *g*, and the lower organic phase was collected and evaporated. Extracted fatty acids were methylated using a Fatty Acid Methylation Kit (Nacalai Tesque, Kyoto, Japan) and purified using a Fatty Acid Methyl Ester Purification Kit (Nacalai Tesque). The eluted solution was evaporated to dryness and kept at –20°C. The residues were solubilized in hexane, and fatty acids were identified by gas chromatography-mass spectroscopy (GC-MS) (JMS-T100 GCV; JEOL, Tokyo, Japan). SCD1 activity was estimated as the oleate-to-stearate ratio (18:1/18:0) and palmitoleate-to-palmitate ratio (16:1/16:0). The 16:1/16:0 ratio seems to be a better indicator of endogenous SCD1 activity than the 18:1/18:0 ratio [23]. The *de novo* lipogenesis index was calculated from the palmitic-to-linoleic acid ratio (16:0/18:2n-6) [24, 25].

### Immunohistochemistry

The mediobasal hypothalamus was injected with AAV-CTL or AAV-NPGL, as described above. Four weeks later, the brain tissues were sectioned into 20-µm-thick slices with a cryostat at −20°C following cryoprotection and freezing. The sections were incubated in blocking buffer (1% bovine serum albumin, 1% normal donkey serum, and 0.3% Triton X-100 in 10 mM phosphate-buffered saline) for 1 h at room temperature before incubation with a rabbit antibody against NPGL (1:100 dilution in blocking buffer), overnight at 4°C. Cy3-conjugated donkey anti-rabbit IgG (1:500 dilution, 711-165-152; Jackson ImmunoResearch Laboratories, West Grove, PA) was used as the secondary antibody. Immunoreactive labeling was observed using a microscope (Eclipse E600; Nikon, Tokyo, Japan).

### Hematoxylin and Eosin Staining

The iWAT was soaked in 4% paraformaldehyde at the endpoint of *Npgl* overexpression, embedded in paraffin, and sectioned to a thickness of 8 µm with a microtome. The sections were then air-dried and deparaffinized in a graded alcohol series. The nucleus and cytoplasm were stained with hematoxylin and eosin (5 min for each stain), and the sections were washed in tap water. After dehydration in a graded alcohol series and clearing with xylene, the sections were mounted on slides and examined under a microscope.

### Oil Red O Staining

To detect fat accumulation in the liver, the liver was fixed in 4% paraformaldehyde and sliced into 10-μm-thick sections. Sections were air-dried, rinsed with 60% isopropanol, stained with Oil Red O solution for 15 min at 37°C, and rinsed with 60% isopropanol. Nuclei were counterstained with hematoxylin for 5 min, and sections were then washed in tap water. Coverslips were applied using an aqueous mounting medium, and microscopic examination was performed using a microscope.

### Serum Biochemical Analysis

Serum levels of glucose, lipids, and insulin were measured using appropriate equipment, reagents, and kits. The GLUCOCARD G+ meter was used to measure glucose content (Arkray, Kyoto, Japan). The NEFA C-Test Wako (Wako Pure Chemical Industries, Osaka, Japan) was used to measure free fatty acid levels. Triglyceride E-Test Wako (Wako Pure Chemical Industries) was used to measure triglyceride levels and the Cholesterol E-Test Wako (Wako Pure Chemical Industries) was used to assess cholesterol content. The Rebis Insulin-mouse T ELISA kit (Shibayagi, Gunma, Japan) was used to measure insulin levels.

### Statistical Analysis

Group differences between AAV-NPGL and AAV-CTL injected animals were assessed using Student’s *t-*test or Welch’s *t-*test. *P* values of < 0.05 were considered significant.

## Results

### Effects of NPGL-precursor Gene Overexpression on Food Intake, Body Mass, and Food Efficiency

AAV-induced overexpression in the mediobasal hypothalamus was confirmed by immunohistochemistry (Supplementary Fig. S1b). Chronic (9 week) *Npgl* overexpression in the hypothalamus significantly increased cumulative food intake and body mass in mice fed NC and a HCD relative to controls (Fig. 1a–d).

**Fig. 1.**
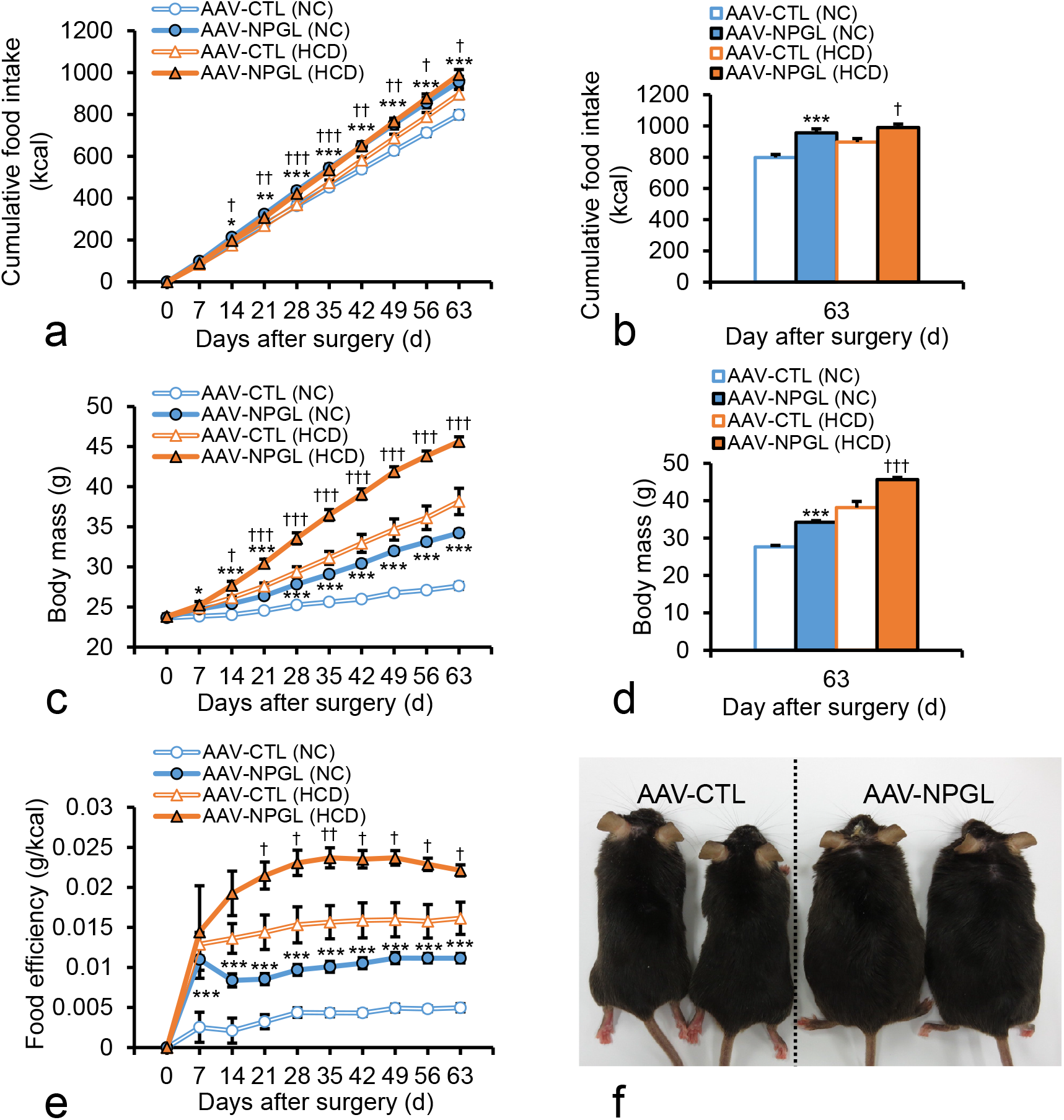
Effects of *Npgl* overexpression on food intake, body mass, and food efficiency. The panels show the data obtained by injection of the AAV-CTL or the AAV-NPGL in mice fed NC or a HCD for 9 weeks. **a** Cumulative food intake at all points. **b** Cumulative food intake at 9 weeks after injection. **c** Body mass at all points. **d** Body mass 9 weeks after injection. **e** Food efficiency is expressed as body weight gain per cumulative food intake per week. **f** Representative photograph of mice at 8 weeks after injection of the AAV-CTL or the AAV-NPGL under a HCD. Each value represents the mean ± standard error of the mean (n = 8/group). \**P* < 0.05, \*\**P* < 0.01, \*\*\**P* < 0.005 vs. AAV-CTL (NC), †*P* < 0.05, ††*P* < 0.01, †††*P* < 0.005 vs. AAV-CTL (HCD). NPGL, neurosecretory protein GL; AAV-CTL, AAV-based control vector; AAV-NPGL, AAV-based NPGL-precursor gene vector; NC, normal chow; HCD, high-calorie diet.

To determine whether the increase in body mass was caused by food intake, we next calculated food efficiency, which is a measure of how much mass is gained per unit of food intake [20]. Food efficiency was markedly increased by *Npgl* overexpression in mice fed NC and those fed a HCD (Fig. 1e). *Npgl*-overexpressing mice fed a HCD were visibly obese, as compared to the AAV-based control vector-injected mice (Fig. 1f).

### Effects of NPGL-precursor Gene Overexpression on Body Composition and Serum Parameters

To address the cause of *Npgl* overexpression-induced body mass increase, we measured the masses of adipose tissues, muscle, and several organs (Figs. 2 and 3). In mice fed NC, the masses of WAT and interscapular brown adipose tissue (BAT) were significantly higher in *Npgl-*overexpressing mice relative to control mice (Fig. 2a, d). Under a HCD, WAT mass and adipocyte size were considerably increased in *Npgl*-overexpressing mice (Fig. 2a–c), whereas there was no difference in the BAT mass in *Npgl*-overexpressing mice (Fig. 2d).

**Fig. 2.**
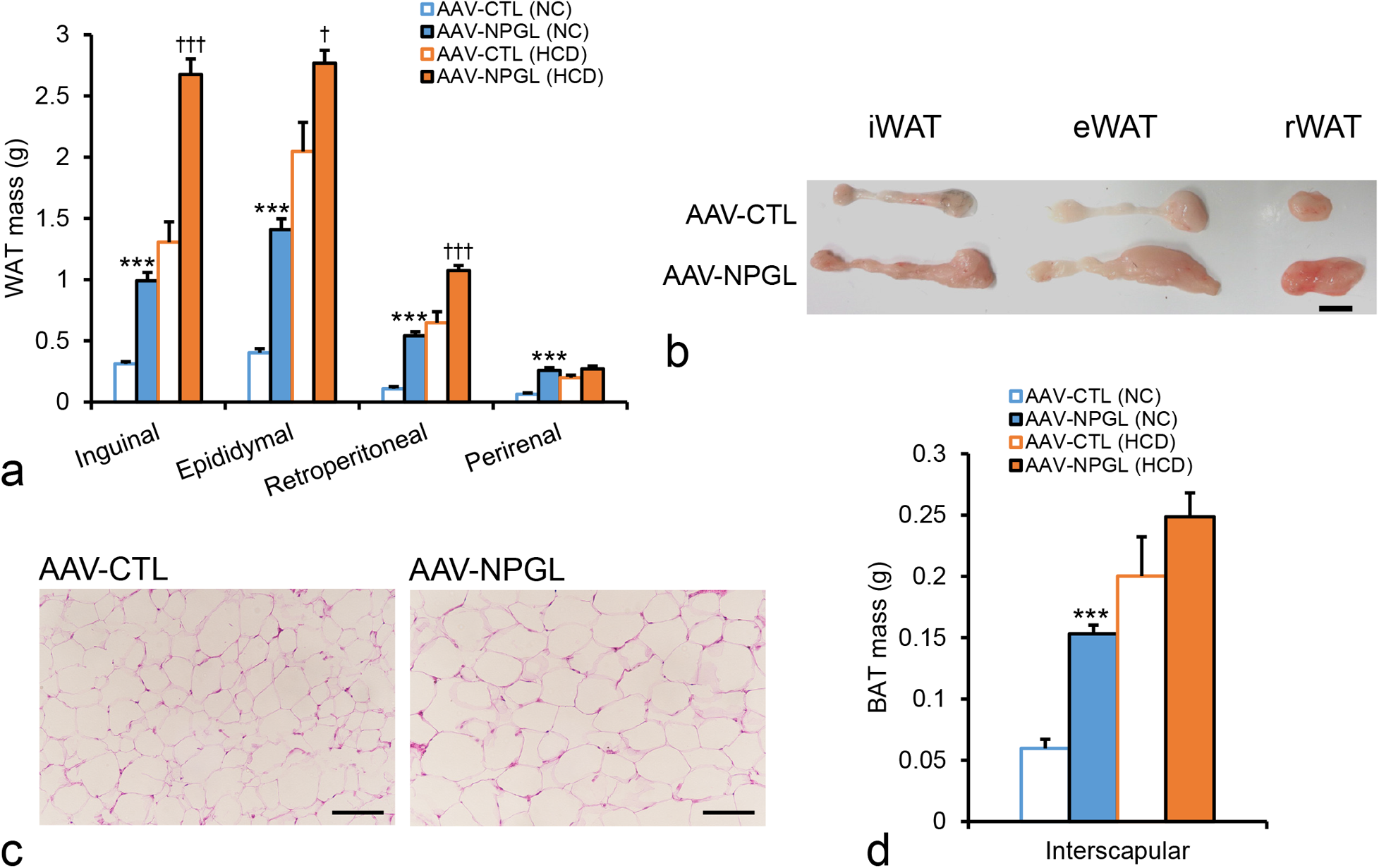
Effects of *Npgl* overexpression on fat accumulation. The panels show the data obtained by injection of the AAV-CTL or the AAV-NPGL in mice fed NC or a HCD for 9 weeks. **a** Mass of the inguinal, epididymal, retroperitoneal, and perirenal WAT. **b** Representative photograph of the iWAT, eWAT, and rWAT in mice fed a HCD. Scale bar = 1 cm. **c** Representative photographs in sections of the iWAT in mice fed a HCD. Scale bar = 100 µm. **d** Mass of the interscapular BAT. Each value represents the mean ± standard error of the mean (n = 8). \*\*\**P* < 0.005 vs. AAV-CTL (NC), †*P* < 0.05, †††*P* < 0.005 vs. AAV-CTL (HCD). NPGL, neurosecretory protein GL; AAV-CTL, AAV-based control vector; AAV-NPGL, AAV-based NPGL-precursor gene vector; NC, normal chow; HCD, high-calorie diet; WAT, white adipose tissue; iWAT, inguinal WAT; eWAT, epididymal WAT; rWAT, retroperitoneal WAT; BAT, brown adipose tissue.

**Fig. 3.**
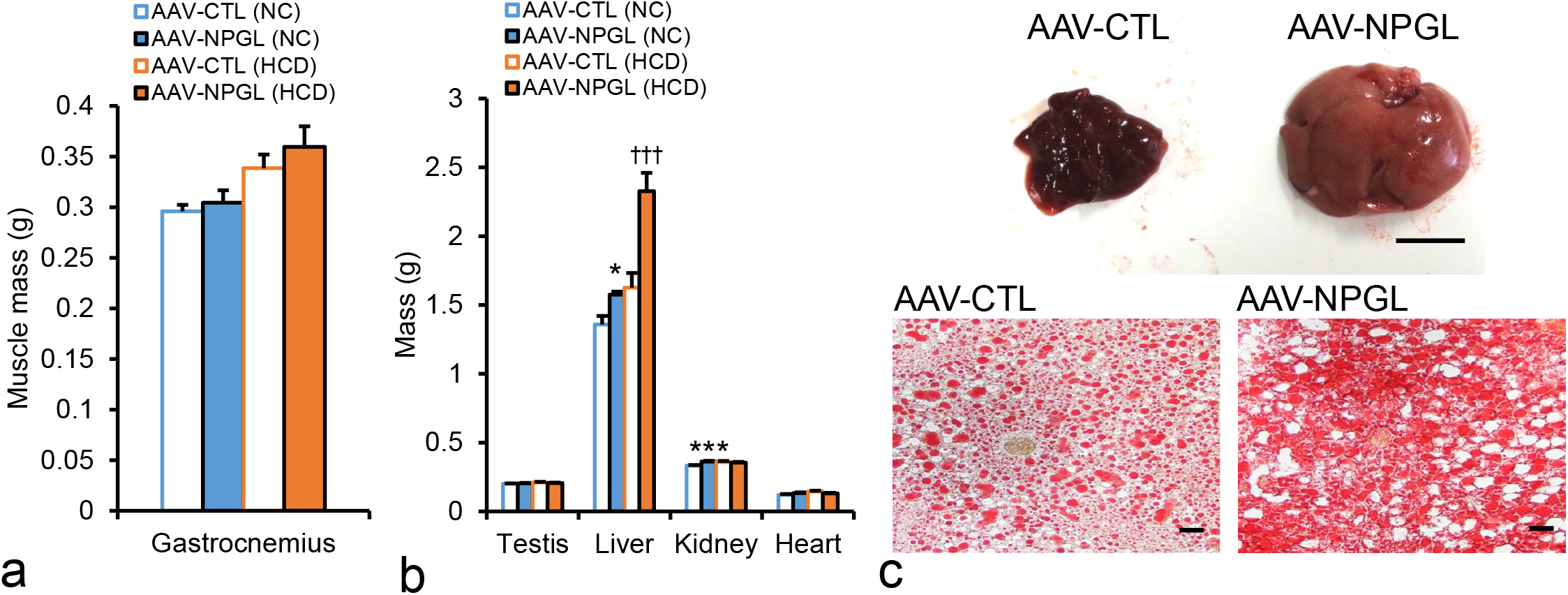
Effects of *Npgl* overexpression on the muscle and organs. The panels show the data obtained by injection of the AAV-CTL or the AAV-NPGL in mice fed NC or a HCD for 9 weeks. **a** Mass of the gastrocnemius muscle. **b** Mass of the testis, liver, kidney, and heart. **c** Representative photograph of the liver and liver sections stained by Oil Red O in mice fed a HCD. Scale bars = 1 cm in the top and 100 µm in the bottom. Each value represents the mean ± standard error of the mean (n = 8). \**P* < 0.05, \*\*\**P* < 0.005 vs. AAV-CTL (NC), †††*P* < 0.005 vs. AAV-CTL (HCD). NPGL, neurosecretory protein GL; AAV-CTL, AAV-based control vector; AAV-NPGL, AAV-based NPGL-precursor gene vector; NC, normal chow; HCD, high-calorie diet.

While the mass of WAT increased markedly, the mass of the gastrocnemius muscle was not different in *Npgl-*overexpressing mice in either diet condition (Fig. 3a). With regard to peripheral organs, the mass of the liver was increased relative to controls by *Npgl* overexpression under both feeding conditions (Fig. 3b). In addition, *Npgl* overexpression induced a pronounced whitening of the liver in mice fed a HCD (Fig. 3c). To explore the underlying cause for the whitening of the liver, we performed a histological examination of lipid deposition in the liver. Oil Red O staining revealed that lipid droplets were increased in the liver in mice with *Npgl* overexpression (Fig. 3c). The masses of the testis and heart were not changed by *Npgl* overexpression (Fig. 3b). The mass of the kidney increased in *Npgl*-overexpressing mice fed NC, whereas there was no effect when these mice were fed a HCD (Fig. 3b).

Fat accumulation can cause hyperglycemia and hyperlipidemia [5]. To reveal the effects of *Npgl* overexpression on blood parameters, we measured serum levels of glucose, insulin, and lipids. *Npgl* overexpression increased serum levels of insulin and cholesterol under both feeding conditions, whereas serum levels of glucose and free fatty acids were unaffected (Fig. 4a, b, d, e). Moreover, the serum triglyceride level was increased by *Npgl* overexpression in mice fed NC (Fig. 4c).

**Fig. 4.**
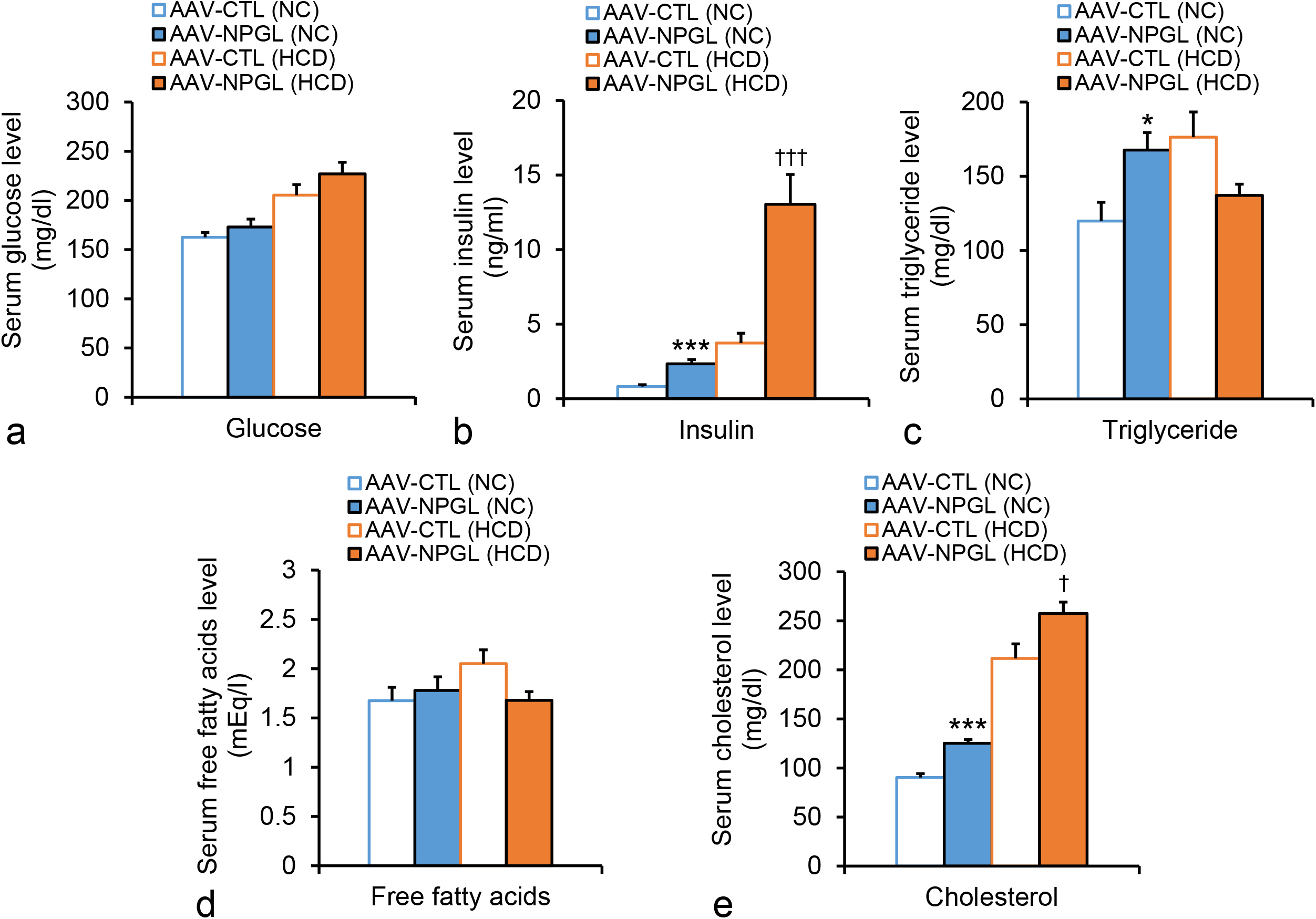
Effects of *Npgl* overexpression on serum parameters. The panels show the data obtained by injection of the AAV-CTL or the AAV-NPGL in mice fed NC or a HCD for 9 weeks. **a** Serum level of glucose. **b** Serum level of insulin. **c** Serum level of triglyceride. **d** Serum level of free fatty acids. **e** Serum level of cholesterol. Each value represents the mean ± standard error of the mean (n = 7–8). \**P* < 0.05, \*\*\**P* < 0.005 vs. AAV-CTL (NC), †*P* < 0.05, †††*P* < 0.005 vs. AAV-CTL (HCD). NPGL, neurosecretory protein GL; AAV-CTL, AAV-based control vector; AAV-NPGL, AAV-based NPGL-precursor gene vector; NC, normal chow; HCD, high-calorie diet.

### Effects of NPGL-precursor Gene Overexpression on mRNA Expression and Endogenous Activity of Lipid Metabolism-related Genes

We detected fat accumulation in the WAT and liver as a result of *Npgl* overexpression, so we analyzed the mRNA expression of lipid metabolism-related genes by quantitative RT-PCR in the iWAT and the livers of mice fed NC or a HCD. The following genes were analyzed: acetyl-CoA carboxylase (*Acc*), fatty acid synthase (*Fas*), *Scd1*, and glycerol-3-phosphate acyltransferase 1 (*Gpat1*) as genes encoding lipogenic enzymes; carbohydrate-responsive element-binding protein α (*Chrebpα*) as a lipogenic transcription factor; carnitine palmitoyltransferase 1a (*Cpt1a*), adipose triglyceride lipase (*Atgl*), hormone-sensitive lipase (*Hsl*), and fibroblast growth factor 21 (*Fgf21*) as genes encoding lipolytic enzymes; glyceraldehyde-3-phosphate dehydrogenase (*Gapdh*) as a carbohydrate metabolism enzyme gene; solute carrier family 2 member 4 and 2 (*Slc2a4, Slc2a2*) as glucose transporters; cluster of differentiation 36 (*Cd36*) as a fatty acid transporter; peroxisome proliferator-activated receptor α (*Pparα*) and γ (*Pparγ*) as genes encoding lipid-activated transcription factors; peroxisome proliferator-activated receptor γ coactivator 1α (*Pgc1α*) as a thermogenic regulator gene; tumor necrosis factor α (*Tnfα*) as an inflammatory cytokine gene; and adiponectin (*Adipoq*) as an anti-inflammatory agent-encoding gene. Under NC condition, mRNA expression levels of *Gpat1, Chrebpα, Cpt1a, Atgl, Hsl, Slc2a4, Cd36, Pparγ*, and *Adipoq* were increased by *Npgl* overexpression in the iWAT (Fig. 5a). In contrast, the mRNA expression level of *Tnfα* was decreased (Fig. 5a). Under a HCD with *Npgl* overexpression, mRNA expression levels of *Chrebpα* were upregulated and those of *Tnfα* were downregulated relative to controls, similar to the observations in mice fed NC with *Npgl* overexpression (Fig. 5a).

**Fig. 5.**
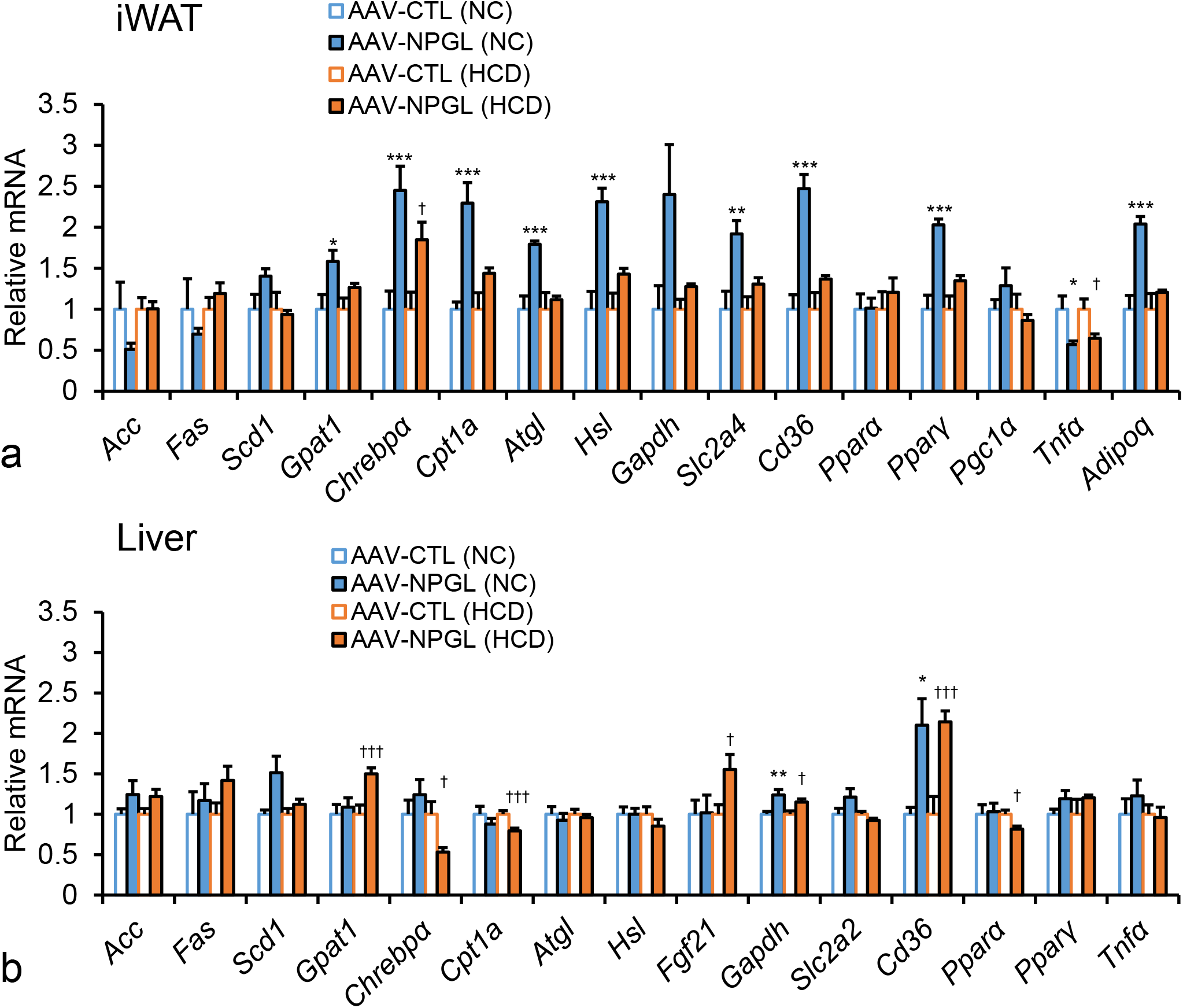
Effects of *Npgl* overexpression on the mRNA expression of lipid metabolism-related genes. The panels show the data obtained by injection of the AAV-CTL or the AAV-NPGL in mice fed NC or a HCD for 9 weeks. **a** mRNA expression levels in the iWAT. **b** mRNA expression levels in the liver. Each value represents the mean ± standard error of the mean (n = 8). \**P* < 0.05, \*\**P* < 0.01, \*\*\**P* < 0.005 vs. AAV-CTL (NC), †*P* < 0.05, †††*P* < 0.005 vs. AAV-CTL (HCD). NPGL, neurosecretory protein GL; AAV-CTL, AAV-based control vector; AAV-NPGL, AAV-based NPGL-precursor gene vector; NC, normal chow; HCD, high-calorie diet; WAT, white adipose tissue; iWAT, inguinal WAT.

In the liver, mRNA expression levels of *Gapdh* and *Cd36* were upregulated in *Npgl-*overexpressing mice fed NC. Under a HCD, mRNA expression levels of *Gpat1, Fgf21, Gapdh*, and *Cd36* were increased by *Npgl* overexpression. In contrast, mRNA expression levels of *Chrebpα, Cpt1a*, and *Pparα* were decreased (Fig. 5b).

To confirm the activity of lipogenic factors in the iWAT, we measured the fatty acid ratio using GC-MS. The fatty acid ratios of palmitoleate-to-palmitate (16:1/16:0) and of oleate-to-stearate (18:1/18:0) indicate increased enzymatic activity of SCD1 [23]. The ratio 16:0/18:2n-6 is an index of *de novo* lipogenesis [24, 25]. The fatty acid ratio of 16:1/16:0 was significantly increased in the iWAT of *Npgl*-overexpressing mice fed NC, although the 18:1/18:0 and 16:0/18:2n-6 ratios remained unchanged (Fig. 6a–c). This result indicates activation of SCD1 by *Npgl* overexpression in the iWAT under NC condition. Under the HCD condition, these ratios did not change in the iWAT upon *Npgl* overexpression (Fig. 6a–c).

**Fig. 6.**
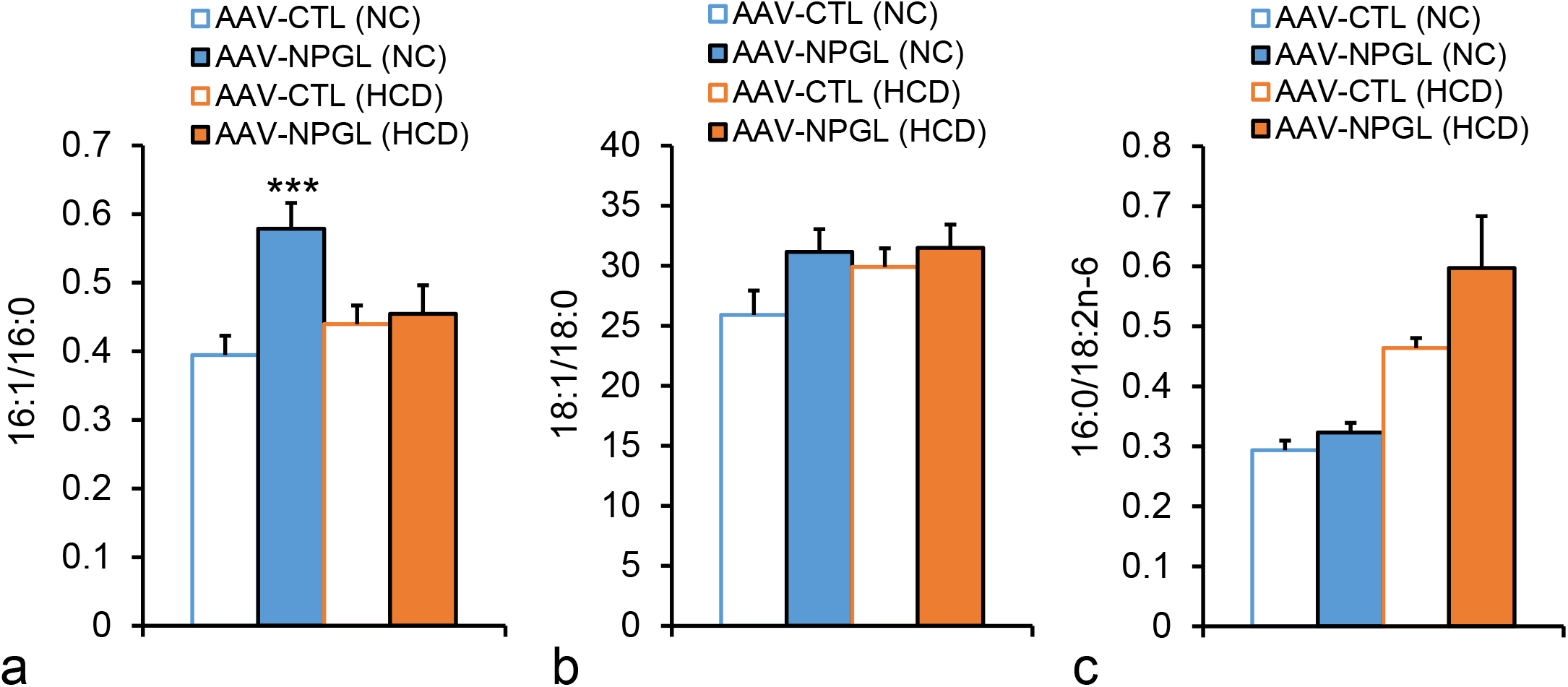
Effects of *Npgl* overexpression on fatty acid ratio in the iWAT. The panels show the data obtained by injection of the AAV-CTL or the AAV-NPGL in mice fed NC or a HCD for 9 weeks. **a** Ratio of fatty acids (16:1/16:0). **b** Ratio of fatty acids (18:1/18:0). **c** Ratio of fatty acids (16:0/18:2n-6). Each value represents the mean ± standard error of the mean (n = 8). \*\*\**P* < 0.005 vs. AAV-CTL (NC). NPGL, neurosecretory protein GL; AAV-CTL, AAV-based control vector; AAV-NPGL, AAV-based NPGL-precursor gene vector; NC, normal chow; HCD, high-calorie diet; WAT, white adipose tissue; iWAT, inguinal WAT.

## Discussion

We have demonstrated that NPGL is involved in energy homeostasis in vertebrates, including birds and rodents [13–16, 26, 27]. In particular, we recently showed that *Npgl* overexpression modestly stimulated feeding behavior and fat accumulation in rats [14]. However, whether NPGL causes obesity across species has not previously been clearly elucidated. In the present study, we found that *Npgl* overexpression in mice induced a remarkable increase in food intake, body mass, and fat mass under both NC and HCD conditions. Therefore, these data indicate that NPGL stimulates fat accumulation in WAT very rapidly, resulting in obesity within 8 weeks in mice.

This difference in the effects of *Npgl* overexpression on body mass change between mice and rats might be a result of species differences in metabolism. Wistar rats were used in the previous study [14], while C57BL/6J mice, which are susceptible to obesity [28], were used in this study. Another possibility may be the difference in the amount of food intake between rats and mice. Although cumulative food intake was slightly increased at the end point in rats fed NC [14], feeding was markedly increased in mice from the early during the treatment period and for the remainder of this experiment in mice (Fig. 1a, b). Using morphological analysis, we previously found that NPGL-containing fibers innervate neurons expressing POMC, a precursor of potent anorexigenic α-MSH, in the mouse hypothalamus [15]. However, this finding was not observed in rats (unpublished observation). Thus, the orexigenic activity of NPGL due to suppression of the activity of POMC neurons in mice is likely higher than that in rats. Future electrophysiological studies are needed to elucidate the efficacy of NPGL in terms of negative modulation of POMC neurons in mice.

The present study revealed that food efficiency was increased by *Npgl* overexpression (Fig. 1e). This result suggests that NPGL-induced fat accumulation is caused not only by energy intake, but also by other factors. Our previous study showed that chronic i.c.v. infusion of NPGL for 13 days reduced energy expenditure and locomotor activity in mice [16]. On the other hand, many studies have demonstrated that obesity resulting from fat accumulation is closely related to inflammation in the WAT [17, 29, 30]. Inflammation in the WAT is mainly regulated by TNFα, an inflammatory cytokine and adiponectin, an anti-inflammatory cytokine [29]. TNFα directly induces lipolysis in adipocytes [31]. Moreover, adiponectin improves insulin sensitivity in the WAT [32]. The present study demonstrated that the mRNA expression of *Tnfα* was decreased, while that of *Adipoq* was increased in iWAT following *Npgl* overexpression (Fig. 5a). Taken together, these results suggest that NPGL suppresses inflammation in adipose tissues, resulting in improved insulin sensitivity and fat accumulation. Further studies are required to clarify the involvement of NPGL in the inflammatory response in the adipose tissue and insulin sensitivity.

Quantitative RT-PCR showed that *Npgl* overexpression increased mRNA expression of factors involved in lipid metabolism in iWAT, especially under the NC condition (Fig. 5a). This result is not surprising as previous findings establish that lipid metabolism is responsive to nutritional conditions [33]. Docosahexaenoic acid, one of the polyunsaturated fatty acids, upregulates lipid metabolic factors, including *Hsl* and *Pparγ* [34]. Specifically, *Hsl* and *Pparγ* mRNA expression were upregulated by *Npgl* overexpression in the iWAT of mice fed NC, but were unchanged under the HCD condition (Fig. 5a). Therefore, our study suggests that intake of HCD may mitigate the effects of NPGL on lipid metabolism. In addition, *Npgl* overexpression upregulated mRNA expression for factors involved in both lipogenesis and lipolysis; also activating SCD1 (Fig. 5a, 6). Lipolysis is regulated by several factors, including ATGL and HSL [35]. In particular, HSL is thought to be the rate-limiting enzyme of lipolysis, and the enzyme is activated by phosphorylation [36, 37]. In DIO mice, HSL protein levels were highly similar to those in control lean mice, although the level of HSL phosphorylation was significantly decreased [38]. Furthermore, insulin stimulates the degradation of cAMP through activation of phosphodiesterase-3B (PDE-3B), suppressing HSL activation [37]. Indeed, we observed an increase in serum insulin levels as a result of *Npgl* overexpression in mice (Fig. 4b). Therefore, it is possible that HSL might not be activated by phosphorylation despite the abundance of *Hsl* mRNA. Conversely, the upregulation of mRNA expression related to lipolysis might be a result of feedback control due to fat accumulation. Further studies are needed to elucidate the effects of NPGL on lipolysis and lipogenesis to reveal the mechanism of fat accumulation in these mice.

*Npgl* overexpression increased liver mass in mice under NC and HCD conditions (Fig. 3b). A previous study reported that elevated insulin stimulates fat accumulation in the liver [39]. In agreement with this finding, we observed increases in serum insulin upon *Npgl* overexpression, as described above (Fig. 4b). Moreover, NPY, which is an orexigenic peptide like NPGL, regulates insulin secretion in rodents [40]. Therefore, the present study suggests that NPGL controls insulin secretion and triggers fat accumulation in the liver. In addition, a previous report revealed that fatty liver leads to hyperlipidemia, including hypercholesteremia and hypertriglyceridemia [41]. It is possible that the fatty liver in mice fed NC and HCD was induced by hyperinsulinemia, resulting in hyperlipidemia (Fig. 4b, c, e). Moreover, despite hyperinsulinemia and hyperlipidemia, blood glucose levels were not changed by *Npgl* overexpression (Fig. 4a, b, c, e). Therefore, NPGL may induce “metabolically healthy obesity”, rather than obesity in the DIO model. This hypothesis is supported by the anti-inflammatory effects of *Npgl* overexpression in the WAT, as described above. Fat accumulation in the liver without excessive alcohol consumption is defined as nonalcoholic fatty liver disease (NAFLD) [42]. Most patients with NAFLD only exhibit simple steatosis [42]. Based on our data in mice, NPGL may also induce simple steatosis rather than nonalcoholic steatohepatitis (NASH). Further investigations are needed to clarify the effects of NPGL on insulin secretion, blood glucose homeostasis, and steatosis in the liver.

In conclusion, *Npgl* overexpression in mice enhances feeding behavior, fat accumulation, and secretion of insulin, rapidly resulting in obesity as compared to the DIO animal model that relies on a high-fat diet. Because *Npgl* overexpression maintains steady-state levels of blood glucose even in obesity, it may be a useful tool for creating a novel animal model of obesity compared to models induced by overeating, such as *ob/ob* mice. Hence, progress in research on NPGL will contribute to a new approach in research on obesity.

## Acknowledgements

We are grateful to Shiki Okamoto for discussion, and Takaya Saito and Atsuki Kadota for experimental support.

## Statement of Ethics

All animal experiments were performed according to the Guide for the Care and Use of Laboratory Animals prepared by Hiroshima University (Higashi-Hiroshima, Japan), and these procedures were approved by the Institutional Animal Care and Use Committee of Hiroshima University (permit numbers: C11-2, C13-12, and C13-17).

## Author Contributions

Conceptualization, Y.N. and K.U.; methodology, Y.N., K.F., K.S., E.I.-U., and M.F.; investigation, Y.N., K.F., K.S., E.I.-U., M.F., G.E.B., L.J.K, and K.U.; writing—original draft preparation, Y.N., and K.F.; writing—review and editing, Y.N., K.F., G.E.B., L.J.K., and K.U.; visualization, Y.N.; project administration, K.U.; funding acquisition, K.F., E.I.-U., and K.U. All authors have read and agreed to the published version of the manuscript.

