## Supplementary Figure for "Hypothalamic Overexpression of Neurosecretory Protein GL Leads to Obesity in Mice"

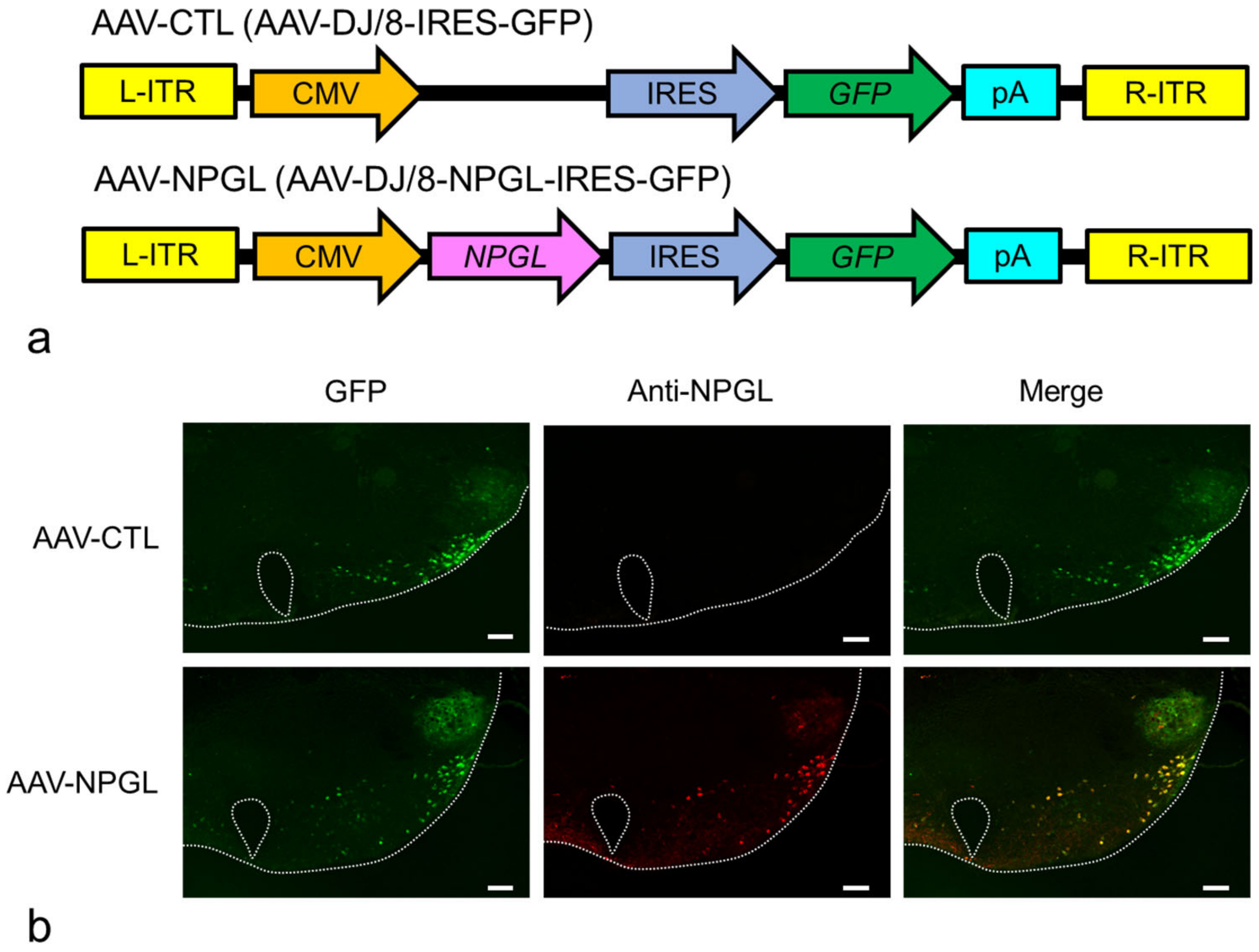


**Supplementary Fig. S1.**
Construction of AAV-based vectors and verification of overexpression. **a** Schematic structures of AAV-CTL and AAV-NPGL. **b** Representative micrograph of the mediobasal hypothalamus at 4 weeks after injection of AAV-CTL or AAV-NPGL. Scale bars = 100 µm. NPGL, neurosecretory protein GL; AAV-CTL, AAV-based control vector; AAV-NPGL, AAV-based NPGL-precursor gene vector; L- and R-ITR, left and right inverted terminal repeat; CMV, cytomegalovirus promoter; IRES, internal ribosome entry site; GFP, green fluorescence protein; pA, polyadenylation signal sequence.
